# Closed-loop automated reaching apparatus (CLARA) for interrogating motor systems

**DOI:** 10.1101/2021.03.01.433419

**Authors:** S Bowles, WR Williamson, D Nettles, J Hickman, CG Welle

**Affiliations:** Department of Neurosurgery, The University of Colorado Anschutz Medical Campus; The NeuroTechnology Center, The University of Colorado Anschutz Medical Campus

## Abstract

*Objective*: Personalized neurostimulation is a rapidly expanding category of therapeutics for a broad range of indications. Development of these innovative neurological devices requires high-throughput systems for closed-loop stimulation of model organisms, while monitoring physiological signals and complex, naturalistic behaviors. To address this need, we developed CLARA, a closed-loop automated reaching apparatus. *Approach:* Using breakthroughs in computer vision, CLARA integrates fully-automated, markerless kinematic tracking of multiple features we use to classify animal behavior and precisely deliver neural stimulation based on behavioral outcomes. CLARA is compatible with advanced neurophysiological tools, enabling the testing of neurostimulation devices and identification of novel neurological biomarkers. *Results*: The CLARA system tracks unconstrained skilled reach behavior in 3D at 150hz without physical markers. The system fully automates trial initiation and pellet delivery and is capable of accurately delivering stimulation in response to trial outcome with sub-quarter second latency. Mice perform the skilled reach task in the CLARA system at a proficiency similar to manually trained animals. Kinematic data from the CLARA system provided novel insights into the dynamics of reach consistency over the course of learning, suggesting that changes are driven entirely by unsuccessful reach accuracy. Additionally, using the closed-loop capabilities of CLARA, we demonstrate that vagus nerve stimulation (VNS) delivered on reach success improves skilled reach performance and increases reach trajectory consistency in healthy animals. *Significance:* The CLARA system is the first mouse behavior apparatus that uses markerless pose tracking to provide real-time closed-loop stimulation in response to the outcome of an unconstrained motor task. Additionally, we demonstrate that the CLARA system was essential for our finding that VNS given after successful completion of a motor task improves performance in healthy animals. This approach has high translational relevance for developing neurostimulation technology based on complex human behavior.

## INTRODUCTION

Personalized neurostimulation therapies, where targeted electrical stimulation is delivered to the nervous system based on an individual’s physiological or behavioral state, have gained prominence in recent years as a potential treatment for numerous nervous system disorders (Olofsson and Tracey, 2017; Peng et al., 2018; Sisterson et al., 2019; Figee and Mayberg, 2021). Traditionally, neurostimulation devices in the clinic have applied open-loop modulation, where stimulation is applied continuously and not altered on the basis of the individual’s state or behavior. These include deep brain stimulation devices for Parkinson’s Disease and Essential Tremor (Benabid et al., 1991, 2009; Kumar et al., 1998; Blomstedt et al., 2010; Cury et al., 2017), Vagus nerve stimulation (VNS) for epilepsy and medically intractable depression (Schachter, 2002; Howland, 2014; Carreno and Frazer, 2017), and spinal cord stimulation (SCS) for pain (Grider et al., 2016; Kapural et al., 2016; Corallo et al., 2020). However, using patient-specific information as feedback to modulate electrical stimulation can optimize therapy and improve patient outcomes (Meidahl et al., 2017; Peng et al., 2018; Swann et al., 2018; Figee and Mayberg, 2021). Patient-specific information required can include neural activity, other physiological metrics such as heart-rate variability, movement, behavior or patient state (Khodaparast et al., 2014; Dawson Jesse et al., 2016; Ciancibello et al., 2019; Sisterson et al., 2019). The important metrics for effective close-looping may depend on the disease or condition. For instance, the Neuropace RNS system for epilepsy uses brain monitoring for epileptiform activity to drive stimulation (Geller et al., 2017; Jobst et al., 2017), while the Inspire Medical neurostimulator for obstructive sleep apnea senses respiration to drive stimulation of the hypoglossal nerve (Maresch, 2018). Recently, investigational studies have explored using movement or behavioral outcomes to provide individualized stimulation control for other disorders. Spinal cord stimulation for the restoration of voluntary movement after injury is improved when stimulation is delivered coincident to the intended movement(Wagner et al., 2018), and vagus nerve stimulation can speed motor rehabilitation following stroke if applied during the movement (Dawson et al., 2016; Engineer et al., 2019). These exciting developments suggest that personalized neurostimulation can optimize therapies for a broad range of indications. To accelerate development of these technologies, new animal model paradigms must be developed that allow for real-time sensing and closed-loop neurostimulation. This will allow for the identification of new physiological or behavioral biomarkers, and the demonstration of the efficacy of closed-loop stimulation. However, the creation of preclinical testing platforms that allow for rapid, responsive neurostimulation poses technical and conceptual challenges that require innovative solutions.

One key challenge to designing these paradigms is the need for real-time measurement of complex behavioral outcomes. This is a critical need, because human rehabilitation is typically focused on complex activities of daily living, rather than highly constrained tasks. State-of-the-art hardware is capable of rapid stimulation delivery, but the ability to collect complex data from the behavior and environment is computationally intensive and can be slow. In addition, rapid classification of the incoming data to determine if a behavioral outcome has been achieved is a challenge. While classification can be achieved manually, automated approaches eliminate experimenter effects and bias (Mah et al., 2021). One approach for rapid, automated data sensing and classification of motor behavior is to rely on measurements from external manipulanda, such as joysticks or levers (Porter et al., 2012; Hays et al., 2014; Hulsey et al., 2016). Yet, these devices constrain behavior in ways that allow animals to become ‘overtrained’ on a very narrow task, and may limit translational relevance to human physical rehabilitation, which focuses more on complex tasks involved in activities of daily living (Dobkin, 2004; Kollen et al., 2006).

Recent advances in computer vision have greatly expanded the approaches that can be used to create closed-loop systems (Mathis et al., 2018). Still, systems using computer vision to date have continued to use basic kinematic thresholds to achieve closed-loop stimulation in real-time (Forys et al., 2020; Sehara et al., 2021), or rely on head-fixation to achieve more uniform input data (Guo et al., 2015; Micallef et al., 2017; Galiñanes et al., 2018). Freely-moving behavior has increased translational relevance, but added complexity in classifying behavioral outcomes. Until now, stimulation in response to complex behavioral outcome has been avoided since it would have to be delivered manually, as is currently the practice in clinical research (Dawson Jesse et al., 2016; Engineer et al., 2019). This increases the labor and time needed to perform experiments, and so systematically testing of stimulation paradigms has been largely relegated to simpler metrics (Porter et al., 2012; Hays et al., 2014; Hulsey et al., 2019). These more complex behaviors also usually come at the cost of gathering detailed quantitative metrics, limiting their analysis of behavioral outcomes to qualitative metrics (Whishaw et al., 2008, 2017).Other systems have succeeded in automation but lack closed-loop stimulation (Williamson et al., 2018; Fenrich et al., 2020).

In addition to automated closed-loop behavioral classification, effective animal model systems must be able to incorporate measures of neural activity and neural manipulations. This provides the capability for identifying new neural biomarkers of effective neurostimulation, and to understand the circuit mechanisms of stimulation to more effectively target stimulation. Thus, systems must be compatible with common neurophysiological tools, including calcium imaging, *in vivo* extracellular recording, and optogenetics. Given these issues in the field there is a clear need for systems that provide quantitative metrics to characterize complex, unconstrained behaviors real-time while remaining compatible with a variety of neurophysiological tools.

In order to address this gap our lab has developed the closed-loop automated reaching apparatus (CLARA) to facilitate multimodal sending and closed-loop stimulation during the learning of a dexterous behavior. CLARA is a fully automated, high-throughput system for training animals to perform a dexterous reaching task while applying closed-loop stimulation based on behavioral outcomes. By using recent advances in computer vision(Mathis et al., 2018), CLARA can track mouse forelimb positional and kinematic information in real time (150 frames/second), and perform classification of complex behavioral outcomes (such as successful pellet retrieval). The system is capable of delivering electrical or optogenetic stimulation based on kinematic features or subjective behavioral outcome. Additionally, CLARA also gathers detailed 3D kinematic data during sessions for quantitative analysis of reaching behavior. Lastly, the kinematic data gathered from CLARA can be combined with simultaneously recorded neural data from Ca2+ imaging or *in vivo* electrophysiology to extract behaviorally-relevant activity without head fixation. In this paper we will describe how advancements in deep learning, novel behavioral devices, and custom software packages were combined to create a data pipeline for efficiently testing closed-loop stimulation in murine models.

## METHODS

### Animal care

All animal procedures were performed in accordance with protocols approved by the Institutional Animal Care and Use Committee at the University of Colorado Anschutz Medical Campus. Male and female adult wild-type C57BL/6 mice between the age of 2 months and one year old were used for all experiments. Mice were group-house prior to surgery and single-housed following surgery and during behavioral experiments. Mice were kept on a 14h light/10h dark cycle with ad libitum access to food and water except during food restriction for behavior training (see Skilled Reach Task).

### Skilled reach task

This task requires a mouse to reach out of a 1 cm wide slit in the front of a behavior box to grab a food pellet with their right paw and retrieve the pellet into the box (Farr and Whishaw, 2002; Whishaw et al., 2008). The pellet is located on a post that is 1 cm away from the front of the box, 1 cm above the cage floor, and 0.5 cm to the left of slit center. A successful reach occurs when a mouse pulls the pellet inside the behavior box using only its right paw (Figure 3A). Mice were food-deprived prior to training to reduce weight to 85-90% of free feeding weight. This weight was checked daily and maintained throughout the training period. The mice were acclimated to the behavior box for two days prior to training. On the first day the researcher fed pellets to the mouse at the slit in the cage. On the second day of acclimation mice were encouraged to reach for post by holding pellets progressively further from the box using forceps. Acclimation ended when mice performed two successful reaches. After acclimation mice were trained on the task for 8 consecutive days, with one 20-minute training session each day (Figure 3A). In VNS implanted mice, mice received stimulation (30Hz, 100us pulse-width, 0.6mA, 0.5s duration) from the CLARA system after successful reaches, defined the mouse bringing a pellet into the behavior box using the right paw. Success rate, the percent of total reach attempts that resulted in a successful pellet retrieval, for each session was determined using real-time stimulation data, and verified by a researcher during post hoc curation of the session videos.

### VNS implant surgery

Mice were anesthetized with 4.5% isoflurane anesthesia and maintained with 1.5% isoflurane. Body temperature was maintained at 37°C using a thermostat-controlled heating pad (TC-1000, CWE Inc.) and received sub-cutaneous lactated ringers (∼100μL/hour). The cervical vagus nerve was accessed through an incision in the ventral cervical region (Figure 5A) and was blunt dissected from the carotid sheath. An additional incision was made at the base of the dorsal skull, and a commercial cuff from Micro-leads (Nano-Fiber Fusion cuff, 150µm internal diameter) was tunneled subcutaneously from the dorsal incision to the ventral incision. The vagus nerve was placed in the cuff, and functional contact was verified by observing a Hering-Breuer reflex, a pause in inspiratory activity (McAllen et al., 2017), in response to cuff stimulation. The ventral incision was sutured using 6-0 absorbable sutures. The skull at the dorsal incision was cleaned using sterile saline and ethanol. The electrical connectors from the cuff were then fixed to the skull using dental cement (C&B Metabond). Skin adjacent to the dental cement was sealed using GLUture (WPI Inc.). Mice received gentamicin (i.m. 3mg/kg), and analgesic meloxicam (i.p. 5mg/kg) following surgery and as needed subsequently. Mice were monitored for 7 days post-surgery to ensure proper recovery.

### In vivo recording electrode implant surgery

Chronic *in vivo* extracellular recordings of single unit activity in motor cortex were conducted during reach training sessions. Mice were anesthetized with 4.5% isoflurane anesthesia and maintained with 1.5% isoflurane. Body temperature was maintained at 37°C using a thermostat-controlled heating pad and received sub-cutaneous lactated ringers (∼100μL as necessary). The dorsal skull was accessed using a vertical incision approx. 1.5 cm long. The skull at the dorsal incision was cleaned using sterile saline and ethanol. A craniotomy was drilled above the forelimb area of M1 (0.3mm anterior, 1.5mm lateral of bregma). Using a stereotaxic device, a single 1.6-mm recording electrode (NeuroNexus) was inserted into the cortex. A stainless steel ground wire was inserted into a burr hole over the cerebellum. The electrode was fixed to the skull using dental cement (C&B Metabond) and surrounding skin was sealed using GLUture (WPI Inc.). Mice were monitored for 7 days post-surgery to ensure proper recovery.

### Electrophysiological recording

Data was acquired at 30kHz using Cheetah acquisition software and Digital Lynx SX hardware (Neuralynx). TTL pulses generated by the master camera were channeled into the Digital Lynx to synchronize videos with the electrophysiology data (Figure 6B). After recording, single-unit activity was clustered manually using Spike Sort 3D (Neuralynx). Isolation distance and L-ratio were used to quantify cluster quality and noise contamination(Schmitzer-Torbert et al., 2005). Frames containing the maximum outward extent of a reach were identified using the CLARA curator (Supp. Figure 1). These frames were used to create reaching epochs in the recording data (±500ms from the reach maximum). Spike data during each reaching epoch was binned at 10 ms and trial-averaged. Firing rate was normalized to baseline activity (1000-500ms before Reach max) using a z-score. Units that displayed a significantly increased (z>2.50) firing rate for at least 100ms during the reach epoch (−500ms before Reach max to 500ms after) were classified as “Movement related”. All other units were classified as “non-movement related”. After all units were classified, they were normalized to generate a heatmap. All units were temporally shifted so that reach max was t=0 and plotted using custom software in MATLAB (MathWorks).

**Figure 1:**
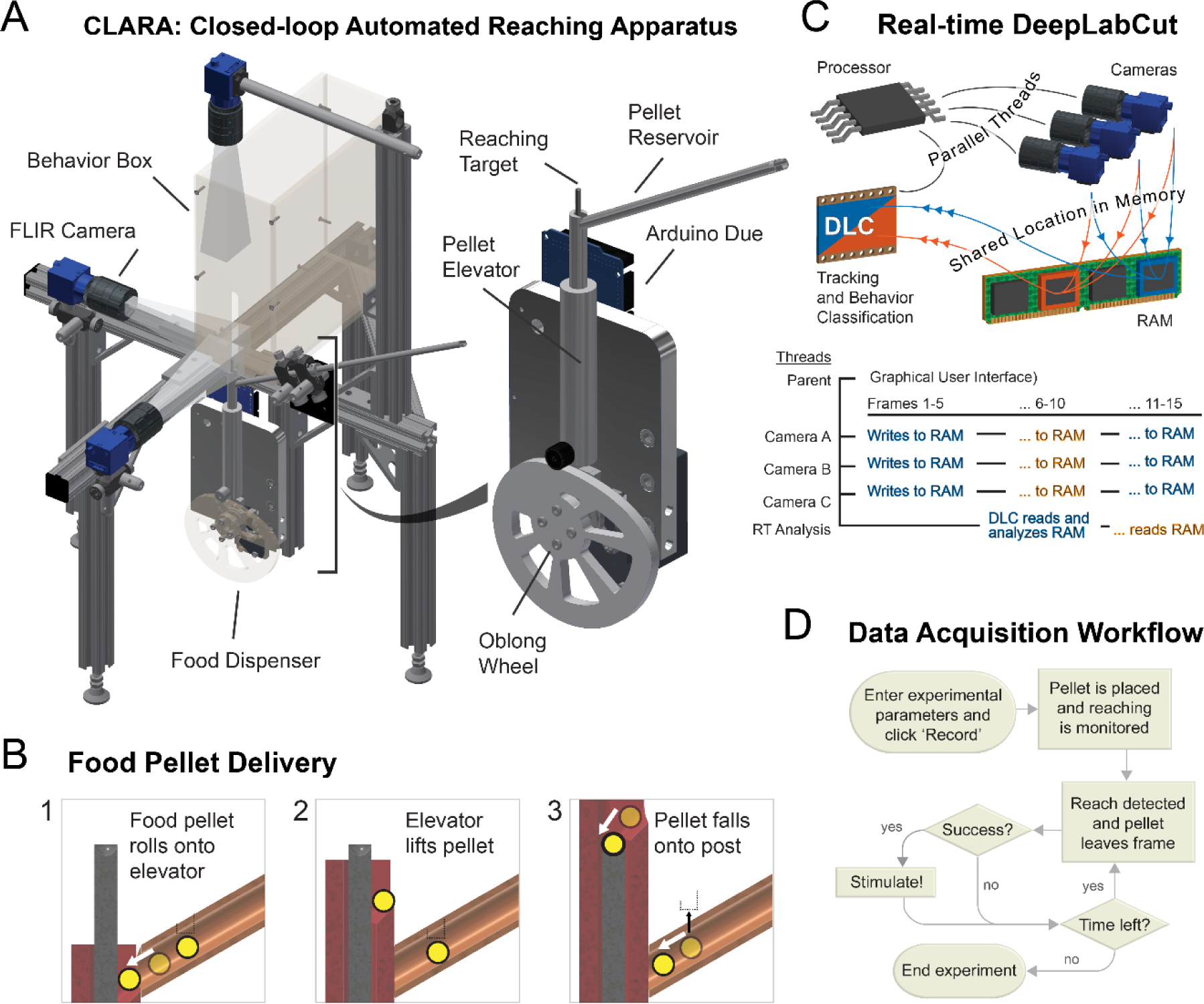
Closed-loop Automated Reaching Apparatus (CLARA). A) Schematic of the CLARA hardware setup. Inset shows custom automated pellet elevator system. B) Workflow for real-time behavior classification. 5 frames from each camera are analyzed by a DLC network simultaneously allowing for real-time analysis at 30Hz with a 150Hz frame rate. C) Pellet elevator allows for automated initiation of behavior trials. D) Full workflow of real-time data acquisition, all steps are fully automated.

### Data acquisition hardware

Images were acquired at 150 Hz using FLIR Blackfly® S (model BFS-U3-16S2M-CS, Edmund Optics, Barrington, NJ) cameras, configured for synchronous recording according to manufacturer instructions. To achieve the desired field-of-view 5-inches from the target, we used a 16mm C-mount fixed focal length lens and a C-to-CS mount adapter, both from Edmund Optics. One of the cameras obtains X/Y information about the mouse hand, a second obtains Z/Y information, and the third is used to see the pellet in the hand during retrieval. We added 1-watt LEDs at a wavelength of 720nm (ThorLabs) and a 60° dispersing lens to illuminate the experiment, with one LED per camera. Using 1-inch-wide t-slotted framing, we assembled a base structure for mounting all hardware. This ensured consistent relative positioning of the behavior box, cameras, LEDs, and the food pellet. Data acquisition was performed using a Dell Precision 7820 (Dell, Round Rock, TX) Linux workstation with an Ubuntu operating system (version 16.04). On each computer, we installed a GeForce RTX 2080 Ti graphics card for real-time tracking. Due to the high power consumption of the cameras, we had to install a SATA-powered PCIe USB 3.1 card. In order to write 450 FPS in real-time, we installed a 2-TB NVMe (Gen 4) hard drive. The computer communicated to an Arduino Due via serial communication to manage the peripheral electronics of the CLARA system. For additional information regarding machined parts, commercially available hardware, and assembly instructions, see link info. Videos were tracked overnight locally on the computer used to acquire the data, and a Synology NAS was used to manage data in a centralized location. The pellet delivery system was designed to deliver the following product: Dustless Precision Pellets® Rodent, Purified, 20mg, Product # F0071 (Bio-Serv, Flemington, NJ). The custom software used to acquire, analyze, and curate the data, as well as installation instructions can be found <u>here</u>.

### Object tracking

Prior to training a neural network, custom software written in Python was used to manually annotate videos by hand. The software features a side-by-side view of all three camera angles, and the option of skipping ahead to events of interest using event files created by CLARA during video acquisition. For example, the user can jump to the next frame on which a pellet was placed. Separate labels were generated for distinct configurations of the mouse hand: flat, spread, and retrieve. Two categories of pellet labels were generated as well to track the pellet when it is in the mouse hand and when it is not. We found that the ability of the network to identify both the pellet and the mouse hand in general was more accurate using this approach. The labels and video frames were packaged in a way that could be interpreted by DeepLabCut, and the Resnet50 pretrained model was selected to train our neural network for improved performance with real-time tracking. After obtaining at least 100 labels for each category, network training proceeded for 1.03 million iterations to a loss of 0.0004. This network was used for real-time tracking (data not saved) and for post-hoc tracking (data saved for subsequent analysis). By design, when DeepLabCut analyzes a video, each video frame receives a best-guess position (x and y) and a confidence value (p) for every label originally used to train the network. In our system, a mouse hand is considered positively identified in the frame if any of the hand categories exceed an empirically determined confidence threshold. Then, the category having the highest confidence score provides information about the state of the hand (flat, spread, or retrieve). In a similar manner, if any of the food pellet categories exceed the confidence threshold, we not only learn where the pellet is, but whether it is in the mouse hand or not. This information greatly improved the accuracy of our behavior classifier.

### Data management

Raw data folders are named with the following hierarchy: date (yyyymmdd), CLARA rig reference (we have 6), and session number, an incremental number starting at 001. At a set time every night, the Linux scheduler, cron, executes a custom Python function which tracks any newly acquired videos locally on the CLARA computer and copies all files (compressed videos, tracking data, metadata, timestamps, and the event file) to the central Synology NAS. Then, cron executes a custom MATLAB function that packages the tracking data into a MATLAB-friendly file format and analyzes the data from each session to approximate the beginning, middle, and end of all reach events, saving the results in a tab-delimited text file. When using the curation GUI to refine the timing of the reach-related events, an Excel file is generated locally on the computer of the user. This provides easy access to their data and allows them to organize their data according to their needs.

### Kinematic analysis

The reach-related event information was used to extract reach epochs from the tracking data. To compare multiple reach trajectories, they were interpolated using dynamic time warping; the full reach trajectory was divided into *n* number of equal length segments independent of velocity(Li et al., 2017). Velocity data was interpolated using the interp3 function in MATLAB. Velocity in the x, y, and z dimension was averaged for each reaching session (1 per day, 20 minute duration), and the location of the peak velocity in each dimension during the reach was identified. Each dimension was compared to determine if peak velocity occurred at different timepoints during the reach. The interpolated reach trajectories were each correlated to every other reach during a session to determine how similar each reach was to the session average. We referred to the resulting average correlation as ‘reach consistency’ and tested within cohorts to determine whether consistency varied over learning or VNS manipulation.

### Statistical tests

The following statistical tests were conducted using GraphPad Prism and all statistical tests are specified in the figure legend. Longitudinal data gathered for velocity was compared using repeated measures ANOVAs. Reaching performance, reach correlation, and reach targeting data from early phases of learning was compared to later timepoints using a paired Student’s t-test. Reach performance and number of reach attempts of researcher and CLARA trained animals were compared using an unpaired Student’s t-test. The improvement of performance and reach correlation was compared to normalized control data using a on sample t-test. Normality tests were conducted using an F test of variance.

## RESULTS

### A closed-loop automated reaching apparatus (CLARA) for neurostimulation based on motor behavior outcome

There is a clear need in the field for closed-loop, automated systems that can classify translationally-relevant complex behavioral outcomes, such as success during motor rehabilitation. Current approaches have been limited by long latency, a lack of automation, behavioral constraints or an inability to integrate with neurophysiological measurements and techniques. We describe a novel system that addresses these challenges, and we demonstrate the ability to modify behavioral outcome through closed-loop neurostimulation.

#### Device description

When creating a behavior system that would allow us to test the underlying mechanisms of electronic medicine, we needed to meet three key criteria: The rig must be automated and high-throughput, the rig must be able to accurately provide closed-loop stimulation based on real-time animal behavior, and the rig must be compatible with a variety of neurophysiological tools. Using these criteria, we developed the Closed-Loop Automated Reaching Apparatus (CLARA) (Figure 1A). CLARA is an apparatus that automatically acquires data for the study of a complex reaching task in freely behaving mice. Here we present three innovations: an automatic pellet delivery system, high-speed object tracking, and a low-latency behavior classifier. The mechanical components include a behavior box, a series of cameras, and a food pellet dispenser (Figures 1A&B, Methods). A single workstation computer manages the cameras, communicates with the microcontroller, and analyzes mouse behavior in real-time using modules extracted from DeepLabCut (Figure 1A&C, (Mathis et al., 2018)). Behavior classification was used for closed-loop stimulation upon the detection of a predetermined mouse behavior, the successful retrieval of a food pellet (Figure 1D).

Pellet delivery is accomplished using a novel system of hardware including an oblong wheel, a sliding elevator, a pellet reservoir, and a narrow post that holds the dispensed pellet at the reaching target (Figure 1A). At the lowered position, a small cavity in the elevator receives a food pellet, and pellet retainer rests on top of a subsequent pellet, holding it in place (Figure 1B, step 1). As the oblong wheel turns, its radius slowly increases, thereby slowly lifting the elevator, which raises the food pellet (Figure 1B, step 2). At its peak, the elevator cavity is slightly above the top of the post, and the pellet rolls onto the post due to gravity and the base of the cavity being angled toward the post. The elevator at this position also raises the pellet retainer to release a subsequent pellet into the cavity (Figure 1B, step 3). The wheel turns to its initial position, and even though the wheel turns at the same speed, the reducing radius quickly reveals the food pellet. The food pellet delivery system was positioned underneath the rest of the hardware to avoid obstructing the camera field of view and to provide access for neural recording technology.

### CLARA allows for high-speed tracking and behavioral classification

High-speed tracking analyzes 200-by-200-pixel regions from each of the FLIR Blackfly® S cameras (model BFS-U3-16S2M-CS, Edmund Optics, Barrington, NJ) acquiring data at 150 frames-per-second (FPS), determining the location of the food pellet and the mouse hand. For tracking to occur in real-time with three cameras recording at 150 FPS, frames must be analyzed at a minimum rate of 450 FPS. This task required creative management of computer memory and processing threads as described in Figure 1C. Our custom data-acquisition software written in Python manages four separate parallel computing threads: one for each camera acquiring frames and one for tracking. It has been previously shown that tracking is more efficient when multiple frames are analyzed in batches (Mathis and Warren, 2018), and we determined that 15 frames (5 per camera) was most efficient on our system. For the sake of computational efficiency, locations in memory are pre-allocated to accommodate the 15 frames of data. However, due to inherent hardware limitations, it is not possible for the cameras to write to memory while the real-time tracking thread reads from the same location in memory. Our solution was to pre-allocate two sets of 15 frames each, and while the cameras are writing frames to one memory-set, the tracking module analyzes the other. When both tasks are complete, all threads are pointed to the other memory-set; the set previously populated by the cameras is tracked, and the cameras populate the set already analyzed. Additional shared locations in memory (the size of a single integer) indicate the status of each thread. In a similar way, the parent thread manages the activity of the child threads, querying the status or sending commands to the cameras or the tracking module.

To classify the behavior, information from the real-time object tracker is used to initiate a trial, and identify features the reaching behavior of the mouse. Multiple markers were tracked simultaneously in a manner that provides positional information as well as behavioral state (see Methods). The primary objects of interest include the pellet and the center of the mouse hand. To initiate a trial, multiple logical decisions are considered. If a pellet is not detected on the post, the elevator is lifted. If the hand of the mouse is inside the box, then the mouse is considered ready to receive a food pellet, and the elevator is dropped to reveal the pellet. After this, tracking information is used to monitor the location of the pellet. If the pellet is no longer on the post, the classifier then determines whether the mouse successfully retrieved the pellet (a “successful reach”) or not (a “failed attempt”). Successful reaches meet the following criteria: reaching behavior is detected, the pellet leaves the post, the mouse hand retracts into the box, and the pellet is never detected away from the mouse hand. Failed attempts occur when the pellet leaves the post and is detected away from the mouse hand. If a reach is categorized as a success, a TTL pulse is delivered from the Arduino board on the CLARA system to trigger stimulation. After a brief wait period of 5 seconds, another pellet is placed. If there is time remaining, this cycle continues until the end of the 20-minute experiment (Figure 1D). Raw data is organized by date, and the following information is saved: the full-length video from each camera, timestamps for each frame, and metadata associated with the experiment. Events that occur during the experiment are also saved, such as the frame on which a pellet was placed or when closed-loop stimulation was delivered.

### CLARA classifies reach outcome with high sensitivity and specificity, producing automated behavioral training comparable to manual training

#### Data curation and post-processing

Every night after data is collected using the CLARA system, a program automatically tracks the videos and copies the data from all CLARA systems to a local server. A separate computer then analyzes the tracking data, identifies reaching events, and records the frames on which reaching begins (Reach Initiation), peaks (Reach Maximum), and either returns to the behavior box or initiates another reach (Reach End, Figure 2A). Another custom GUI written in Python (Supp. Figure 1) allows the user to quickly view and adjust the timing of these events. This GUI also allows the user to assess the accuracy of the behavior classification. Event times were binned relative to the end of the reach and demonstrated that the reach duration from reach initiation and to reach end was 370±120ms. The reach maximum (defined as the maximum extent of the hand away from reach initiation) occurred 221±98ms prior to the reach end (Figure 2B). The behavior classifier consistently identified behavior outcome 224±49ms after reach end, though this is not a measure of system latency, as the behavior classifier runs in real-time, while reach features are identified post-hoc. The latency of the classifier in real time is 151± 32ms after the paw enters the behavior box. The hardware latency between classification and stimulation was less than a single frame (less than 7ms), which is negligible for our purposes (Supp. Figure 2). After curating 12,219 reaches, we determined the classifier had extremely high specificity, with false positive successes classified on 1.12% of all reaches (Figure 2C). The classifier was also highly sensitive, correctly identifying successful outcomes 74.16% of the time resulting in an overall accuracy of 91.17% (Figure 2C). Together this data demonstrates that the CLARA system can deliver closed-loop stimulation based on the classification of a complex, unconstrained behavioral outcome with limited delay and a high degree of specificity.

**Figure 2:**
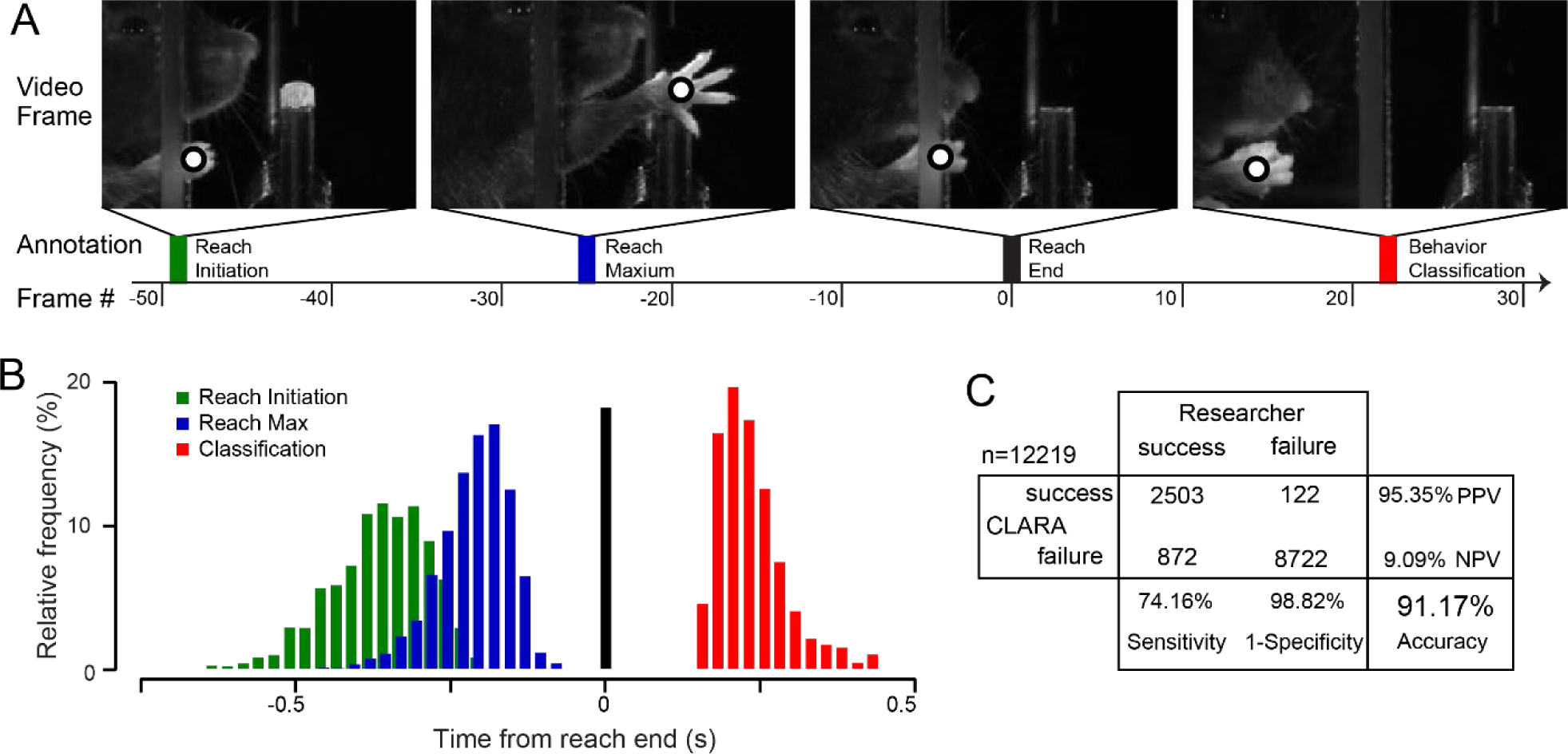
CLARA produces highly accurate real-time classification with sub-second latency. A) Example images of key reach features: Reach initiation (green), maximum outward extension (blue), reach end (black), and behavior classification (red). Frame rate is 150Hz. B) Relative frequency of key features normalized to reach end. reach initiation and maximum occur 370±120ms and 221±98ms ms prior reach end respectively, and classification occurs 224±49ms after reach end. C) Comparison of researcher and CLARA behavior classifications, researcher classification was performed post-hoc.

**Figure 3:**
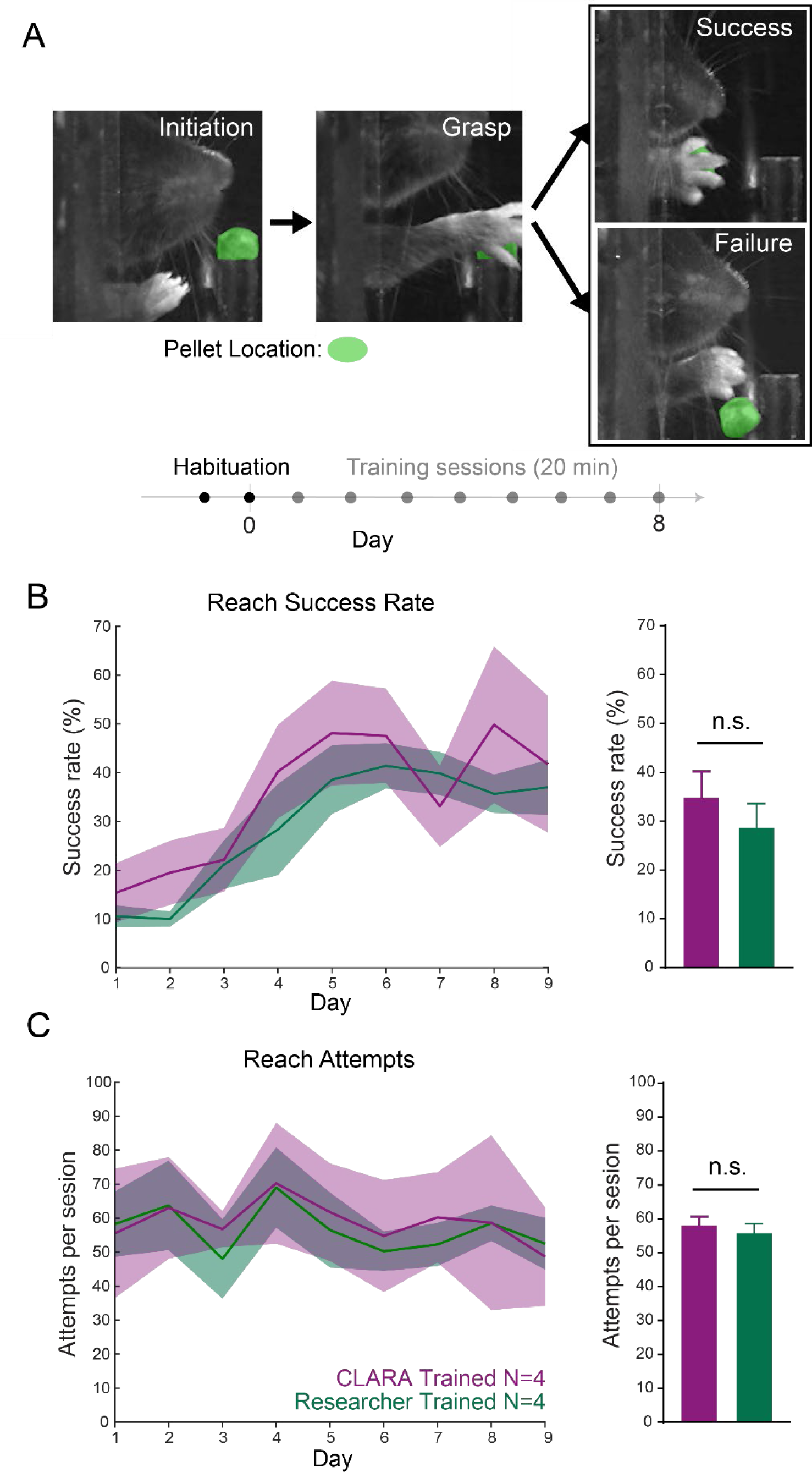
Mice trained in the CLARA system perform similarly to researcher trained controls. A) Example images of a reach and experimental timeline. Successful trials require the pellet to be carried inside the behavior box (top image) while all other results are classified as failures (bottom). Green pseudo-color denotes pellet location. B) Mice trained by the CLARA system (purple) achieve similar rates of success to manually trained animals (green). Bar graph represents average reaching performance per mouse over all days (t=0.999, p=0.3327, unpaired t-test). C) Mice trained by the CLARA system attempt a similar number of reaches to manually trained mice. Bar graph represents average number of attempts per mouse over all days (t=0.758, p=04593, unpaired t-test). Error bars denote standard error.

#### Mouse behavioral performance

The CLARA system is designed to be a high-throughput automated behavioral rig. Mice therefore must be able to learn and perform the skilled reach task in the system with limited researcher supervision. Two cohorts of mice were trained on the skilled reaching task (see Methods). One cohort had food pellets administered manually by a researcher (N=4), while the other cohort had pellets administered automatically through the CLARA system (N=4). A successful reach is defined as the mouse bringing the pellet from the post back into the behavior box using its right paw. The number of successful reaches completed out of the total reaches attempted each session was calculated to determine success rate (Figure 3B). The success rate of the mice was not significantly different between researcher-trained and CLARA-trained animals at any point during training (29.17±12.42 versus 35.32±13.67, p=0.33 unpaired t-test)(Figure 3B). Additionally, the number of total reach attempts made by researcher-trained and CLARA-trained mice was not significantly different (58.84±6.06 versus 56.56±6.72, p=0.46 unpaired t-test)(Figure 3C). From this data we determine that use of the CLARA system does not alter mouse skilled-reaching performance.

### Kinematic tracking with CLARA enables detailed characterization of the reach over learning

#### Detailed 3D Motion Tracking

Skilled reach behavior has been a widely characterized motor behavior paradigm for decades(Gonzalez et al., 2004; Whishaw et al., 2008; Becker and Person, 2019), and is commonly used to model rehabilitation and recovery from neurologic injury(Ryait et al., 2019). However, due to the unconstrained nature of the task, researchers are often limited to qualitative descriptions of the behavior(Whishaw et al., 2008; Mirza Agha et al., 2020). Markerless pose tracking from the CLARA system was used to accurately plot the outward (purple) and return (blue) trajectories of each reach from multiple cameras (Figure 4A). While our quantifications primarily focused on tracking the center of the hand, the system is able to train on nearly any image feature, allowing for high resolution tracking down to the level of individual features (Supp. Figure 3), which can provide other novel features such as grasp dynamics. Using hand tracking data, we extracted several quantitative metrics about each reach, including 3D trajectory and velocity (Figure 4B, pseudocolor represents normalized velocity). This allowed us to observe detailed kinematic features of the reach. For example, outward trajectory of reaches is targeted adjacent to, rather than directly towards, the pellet (Figure 4B). We also observed that velocity changes several times over the course of a reach. By normalizing the length of each reach (see Methods) we were able to measure X, Y, and Z velocity during the outward phase of the reach (Figure 4C). We find that skilled reaching has two distinct phases before pellet contact. Lateral movement towards the pellet (Z velocity) peaks significantly later (55.97±2.16 percent progress between reach initiation and maximum reach extension) in the reach than the max upward (Y velocity, 21.46±0.94 p=0.0001 RM ANOVA) and outward movements (X velocity, 32.61±-3.92 p=0.0001 RM ANOVA) (Figure 4D). Neither the max velocity, nor the mean reach velocity changed over the course of learning (p>0.05, One way ANVOA) (Supp. Figure 4A&B.

**Figure 4:**
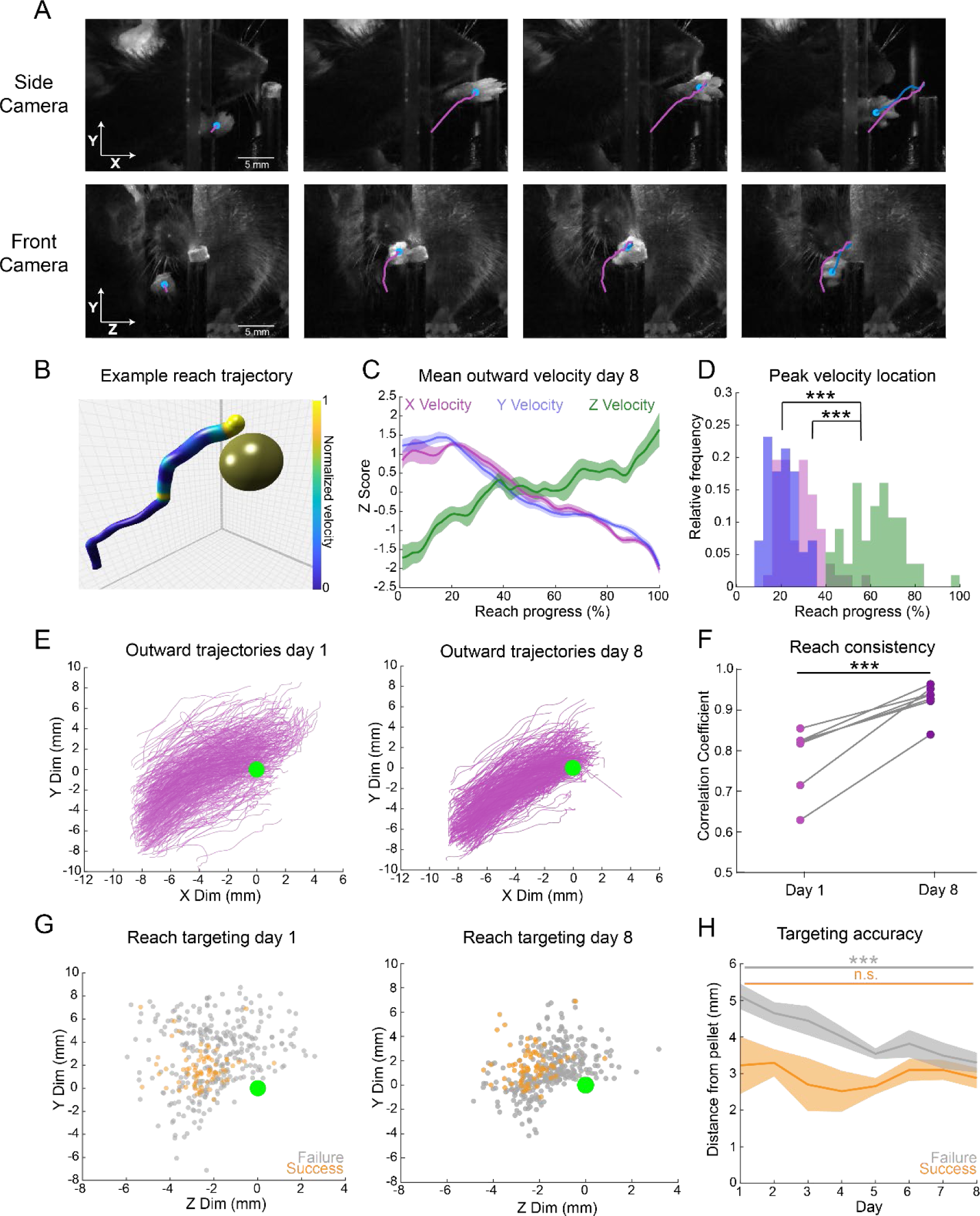
CLARA provides detailed markerless tracking of skilled reach behavior. A) Example frames of a single reach from two camera angles. Purple line denotes outward trajectory and blue line denotes return trajectory, blue dot marks hand center. B) Example reach plotted in 3D space in relation to a pellet (to scale). Colors represent absolute velocity with blue representing high velocity and yellow representing low velocity. C) Normalized outward velocity, plotted as the % completion to reach maximum, on day 8 of reach training in the upward (Y velocity, blue), outward (X velocity, purple) and lateral direction (Z velocity, green). Shaded boxes denote standard error (n=7 mice). D). Distribution of maximum velocity of reaches for each dimension across all days. Max velocity occurs at different points in the outward reach (F=59.15, p=0.0001 RM one-way ANOVA). Time is normalized between reach initiation (0% progress) and maximum reach extension (100% progress). X vs. Z velocity (p=0.0001, Tukey’s HSD) Y vs. Z velocity p=0.0001, Tukey’s HSD). E) Outward reach trajectories from all reaches from the side camera view on day 1 (left) and 8 (right) of training. Green circle represents pellet center (not to scale). F) Mice improve reach consistency during learning. Mean correlation value for all pairs of outward reach trajectories within a session on day 1 and 8 of training, each point represents one animal (n=7) (t=6.350, p=0.0007, paired t-test). G) Reach endpoint (measured from hand center) from n=7 mice at maximum reach extension from the front camera view on day 1 (left) and 8 (right) of training. Orange dots represent successful reaches, grey dots represent reach failures, green circle denotes pellet center. H) Mean absolute distance (mm) of hand center from pellet center at maximum reach extension during learning. Targeting of failure reaches improves from day 1 to 8 (t=7.283, p=0.0003 paired t-test), targeting of successful reaches does not improve (t=1.998, p=0.092 paired t-test.) Shaded boxes represent standard error.

**Figure 5:**
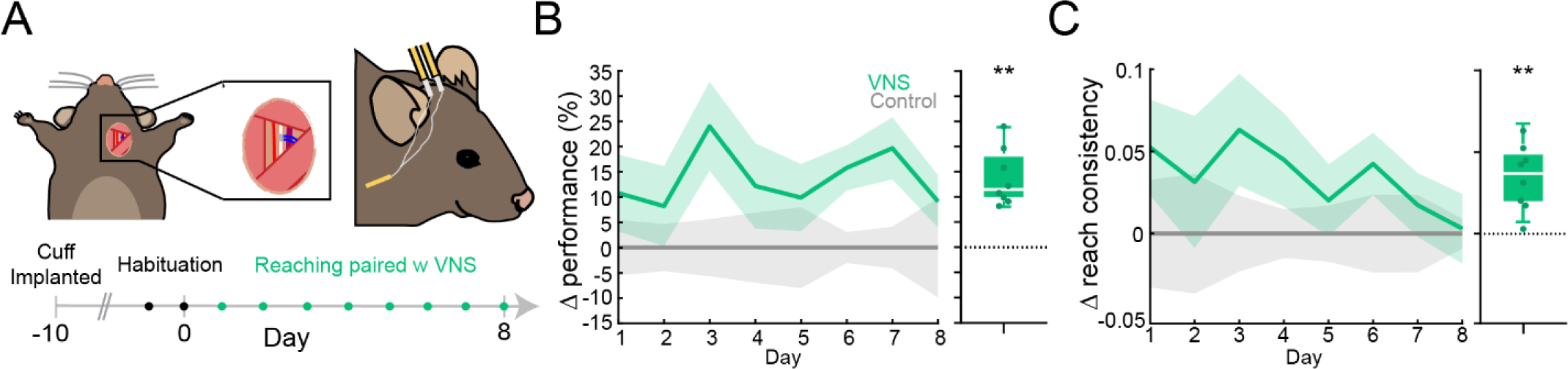
Closed-loop vagus nerve stimulation (VNS) alters skilled reaching behavior in mice. A) Schematic of vague nerve cuff implant surgery and experimental timeline, dots represent one training session. B) Change in task success rate between VNS (green, N=8) and control (grey, n=8) mice across training. Boxplot represent average change in success rate across all days (t=6.11, p=0.0005 one sample t-test). C) Change in reach consistency (mean correlation value for all pairs of reach trajectories) between VNS and control animals. Boxplot represents average change in reach consistency across all days (t=4.85, p=0.019 one sample t-test).

**Figure 6:**
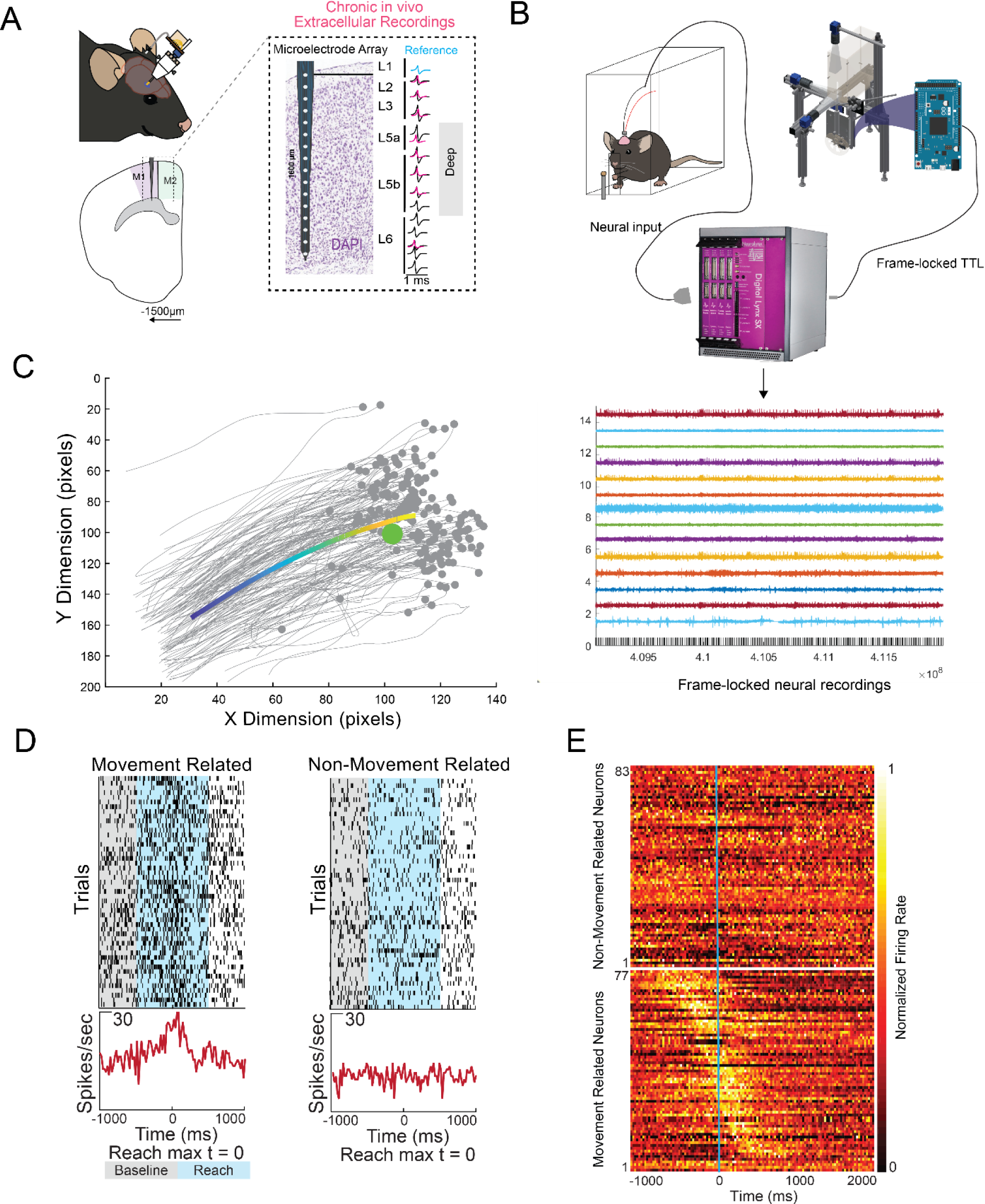
CLARA is Compatible with freely moving neural recording devices. A) Schematic of a chronic extracellular recording electrode implant location. B) CLARA can deliver frame locked TTL triggers to recording hardware (top) to allow neural activity to be matched to individual frames (bottom). Black dashes in bottom image represent frame triggers. C) Example of maximum reach extension timepoints in a trajectory. Grey dots denote max extension, the green circle represents pellet center, and the multicolored line represents mean outward trajectory. D). Example raster plots of a movement related (right) and non-movement related (left) neuron around reach maximum (Time 0). Each trial represents a reach attempt. E) Heatmap of normalized mean firing rate around reach maximum (blue line) for all Non-movement (top) and movement (bottom) related neurons. Dark color denotes low FR while light color denotes high FR.

Previous studies in lever press tasks have shown that reach trajectories become more consistent over learning(Kawai et al., 2015). For our skilled reach task, reach consistency was calculated by temporally warping reaches to normalize their length (see Methods) and then measuring the correlation value between all pairs of outward reach trajectories within a session. In contrast to velocity dynamics, reach consistency changes over eight days of learning, with decreased variability between individual reaches during late learning (day 8) as compared to early learning (day 1) (Figure 4E). The correlation coefficient between all reaches was much higher on the last day of training compared to the first day of training (Mean correlation coefficient 0.78±0.08 on Day1 versus 0.92±0.04 on Day 8, p=0.0007 Paired t-test) (Figure 4F), demonstrating increased consistency.

Increased consistency speaks to the similarity between individual reach trajectories but does not address targeting accuracy. To measure accuracy, we examined the location of the maximum outward hand extension for each reach with respect to the pellet location. As expected, on day 1, successful reach endpoints (orange dots) are closer to the pellet than are failure reach endpoints (grey dots) (Figure 4G). By late learning (day 8), failure endpoints are more tightly clustered. Targeting accuracy increased over time for all reaches (Absolute distance to pellet (mm) 4.66±0.85 Day 1 versus 3.20±0.60 Day 8, p=0.0005 Paired t-test) (Supp. Figure 4C). Improved accuracy was driven by improved targeting of unsuccessful reaches (5.10±0.92 Day 1 versus 3.36±0.72 Day 8, p=0.0003 Paired t-test), as the targeting of successful reaches does not change between day 1 and day 8 (3.56±0.86 Day 1 versus 3.00±0.65 Day 8, p=0.09 Paired t-test) (Figure 4H). The CLARA system provided quantitative kinematic features that allow for characterization of reach, consistency and accuracy over the duration of skilled reach learning.

### The CLARA system demonstrates the effectiveness of precisely-timed VNS to influence motor learning

#### Closed-loop VNS delivery alters reach behavior

The primary purpose of the CLARA system is to determine how closed-loop stimulation paradigms will alter behavior. Previous research has shown that VNS paired with motor rehabilitation can accelerate recovery from movement deficits after stroke (Hays et al., 2014; Khodaparast et al., 2014; Engineer et al., 2019). Yet, there have been no observations of VNS paired with movements influencing movement proficiency in healthy animals, likely due to a lack of precise stimulation during unconstrained behavior. We used the CLARA system to investigate whether VNS could also alter motor learning in healthy animals. To do so, stimulation cuffs were implanted on the cervical vagus nerve in mice(see Methods, (Mughrabi et al., 2020)) and after recovery, the animals were trained to perform the skilled reach task (Figure 5A). The CLARA system delivered stimulation on successful completion of the reach. When VNS was delivered after successful reaches, mice performed the reaching task significantly better than unstimulated controls (11.81±5.465% increase in performance compared to control, p=0.0005 One sample t-test) (Figure 5B). This was accompanied by an increase in the consistency of reach trajectory as measured by the mean correlation value for all pairs (0.035±0.02 increase in mean correlation coefficient compared to control, p=0.0019 One sample t-test) (Figure 6C). This data shows that closed-loop VNS increases skilled reach performance and reduces reach variance in healthy animals when compared to controls. More generally, this demonstrates that CLARA provides accurate and rapid closed-loop stimulation on behavioral outcome, allowing for the exploration of personalized neuromodulation technology.

### CLARA system can be integrated with multimodal physiological measurement methods

#### Integration with multi-channel extracellular recordings

Neurophysiological measurements are needed to identify new biomarkers of disease states and of neurostimulation effectiveness, and also to understanding the neural response to neurostimulation technology. The CLARA system is able to interface with neurophysiological measurement tools of multiple modalities, including extracellular recording, in vivo functional imaging and optogenetic circuit manipulation. Here we demonstrate feasibility of combining CLARA with high-density extracellular recordings of neurons in motor cortex to extract reach-related cortical activity. Extracellular electrode arrays were implanted in the motor cortex of adult mice for chronic recordings. The electrodes spanned the six layers of motor cortex, providing information regarding neural activity throughout the laminar depth (Figure 6A). During a cortical recording, the Arduino on the CLARA system delivered a TTL pulse to the Digital Lynx recording system (NeuraLynx) for each camera frame recorded (Figure 6B). Automatically placed post hoc reach markers (Figure 2A) were then used to identify reach maximum frames, which could be matched to TTL triggers recorded in the session (Figure 6C). This allowed us to separate reach-related and non reach-related units based on their activity during the reach task (Figure 6D). Using the CLARA-generated data, we were able to recapitulate the well described sequential activation of neurons in motor cortex during motor tasks (Figure 6E, (Peters et al., 2014; Adler et al., 2019)). This data demonstrates that CLARA can be used to study *in vivo* neural activity while mice perform freely moving motor tasks. Further, due to the high frame rate detailed tracking, neural activity can be analyzed around specific kinematic features despite the unconstrained nature of the task.

## DISCUSSION

Systematic testing of closed-loop stimulation paradigms for implanted electrical devices in animal models has long been a challenge for the preclinical investigations of personalized neurostimulation devices. The studies require a task that is unconstrained and complex, to replicate typical conditions of human motor rehabilitation (Dobkin, 2004; Engineer et al., 2019). Next, real-time tracking of relevant behavioral variables is needed to perform classification of behavioral features. These classifications then are rapidly trigger neurostimulation, to provide closed-loop stimulation on behaviorally relevant timescales. To optimize stimulation paradigms that best address clinical concerns, these systems need to be automated and capable of high-throughput experiments. The CLARA system is uniquely designed to provide real-time (150 frames/s) tracking and automatic classification of complex, unconstrained behaviors, rapid closed-loop stimulation and compatibility with neurophysiological monitoring tools.

Firstly, CLARA can accurately track and classify the outcome of a complex motor task having high behavioral variability. In other similar behavior systems, the animal interacts with an effector, such as a lever press (Tanaka et al., 2018; Juavinett et al., 2019; Morrison et al., 2019) or joystick (Peters et al., 2017; Bollu et al., 2018). This approach has the advantage of easily extracting positional or force attributes from sensors on the effector. However, these are typically simpler tasks where animals can easily become ‘over-trained’ on the highly stereotyped movements, and animals may not use higher-order cortical motor circuits for the performance of these tasks (Kawai et al., 2015). In contrast, motor cortex is required for forelimb reach task even in highly trained animals (Guo et al., 2015). Our goal was to be able to provide closed-loop stimulation in response to naturalistic behavior, here food retrieval, due to the increased translational relevance to the goal-oriented rehabilitation used in the clinic (Dobkin, 2004). Despite using a more complex and unconstrained task, the CLARA system is still capable of providing closed-loop stimulation with high accuracy and specificity, a key requirement to study certain device applications(Hays et al., 2014).

Secondly, CLARA leverages recent advances in deep learning to allow for markerless pose tracking. Marker-based tracking systems, which rely on the attachment of physical markers to the limbs, are used for closed-loop tracking of naturalistic movements in an unconstrained environment (Peikon et al., 2009; Becker and Person, 2019). These systems directly measure positional data in real-time unlike lever or joystick effector systems. However, the dependency on physical markers limits the flexibility of these systems. In small animal models, affixing multiple markers to a limb can be difficult, particularly at small joints such as fingers. Additionally, closed-loop stimulation still is based on positional or kinematic features (Becker and Person, 2019) which are restricted to where markers are placed. CLARA can instead use information multiple points for tracking, on the resolution of individual digits to provide a wealth of kinematic information.

Multiple studies have implemented markerless tracking of forelimb reach, (Guo et al., 2015; Galiñanes et al., 2018), including some that used DeepLabCut frameworks to provide real-time closed-loop stimulation (Kane et al., 2020; Vonstad et al., 2020; Sehara et al., 2021). However, the majority of these systems involve head fixation, which greatly constrains movement and limits the translational relevance. Further, even in systems that could be used in freely moving contexts (Kane et al., 2020), closed-loop stimulation is limited to positional or positional derivative kinematic thresholds. To our knowledge, CLARA is the first system that can accurately provide close-loop feedback in response to an unconstrained behavior outcome. We were able to achieve this by taking advantage of DeepLabCut’s flexibility in how a network is trained. Prior to training a neural network using DeepLabCut, our custom video-labeling GUI allows the user to add sub-categories to the object being labeled, such as the ‘flat’ hand reaching for a food pellet, a hand ‘spread’ open as it approaches the pellet, or the closed hand attempting to ‘retrieve’ the pellet (see Methods). When the trained network tracks videos using these labels, one gains information about the position of the hand as well as its conformational state related to the behavior of the animal. This allowed simultaneous tracking of the position and state of the hand and pellet in real-time with only a few markers. This combination of state and position was sufficient to distinguish behavior outcomes in the skilled reach task. We believe that by using our application of computer vision in the CLARA system as a model, researchers should be able to rapidly expand the study of behavior driven closed-loop stimulation paradigms in animal models, accelerating the pace of research in the field.

The data we gathered using the CLARA system to study skilled reaching provided a detailed, quantitative description of the reaching task without the use of external markers or time-consuming manual tracking (Card and Dickinson, 2008). We first demonstrated that mice can learn the skilled reach task at a similar level of proficiency in the CLARA system to those trained manually. We then characterized key features of the velocity, trajectory and gross targeting of the reaching behavior, and identified changes to both reach consistency and targeting accuracy over reach learning. Using the closed-loop functionality of the CLARA system, we then demonstrated that a personalized neurostimulation device, VNS, is able to alter motor learning. We found that VNS, when paired with the successful completion of the reach task, improves mice’s ability to perform the task. While VNS has been shown to improve forelimb functional recovery following neurological injury (Porter et al., 2012; Hays et al., 2014; Khodaparast et al., 2014), this is the first study to demonstrate an effect of precisely time VNS on motor function in healthy animals. Moreover, CLARA allowed us to draw new insight into the kinematic changes produced by VNS, demonstrating that VNS increases reach consistency. Reach consistency is a common feature of motor learning that is conserved across model systems (Shmuelof et al., 2012; Kawai et al., 2015; Peters et al., 2017), suggesting that VNS alters a fundamental characteristic of reach learning. These findings demonstrate that the CLARA system can rapidly test if certain neurostimulation paradigms alter motor behavior, and identify how the behavior is changing, giving insight into the underlying mechanism of action.

The CLARA system also has the potential to greatly improve our understanding of the neural effects of electrical stimulation devices. We demonstrate the feasibility of combining CLARA with extracellular neural recordings in motor cortex, and recapitulate the well-characterized sequential activation of motor cortex pyramidal cells during motor movement (Peters et al., 2014; Adler et al., 2019). The flexibility of the tracking system allows for CLARA to be used with a number of neurophysiological tools, such as head mounted miniscopes for in vivo imaging, chronic electrophysiological recording systems and optogenetic stimulation for circuit interrogation.

## CONCLUSION

Future expansion of personalized medicine will require closed-loop capable stimulation devices. High throughput testing of devices and stimulation paradigms will be essential to the development of these devices. Our lab has created the CLARA system, a freely-moving mouse behavioral system that provides automated, closed-loop stimulation based on behavioral outcome by using a novel application of the computer vision framework DeepLabCut. This system provides rapid and accurate neurostimulation to demonstrate the effects of VNS on motor learning in healthy animals. It also provides detailed 3D kinematic data about the skilled reach and is compatible with common *in vivo* neurophysiological tools to explore neural function. This system will provide a model for researchers to rapidly test closed-loop stimulation devices in animal models.

## ACKKNOWLEDGEMENTS

We would like to thank Michael Hall and his machine shop for his delivery of custom parts at high-specifications, as well as Rosenberg Industries in Trimont, MN for their fast, reliable 3D printing services.

---

**Supplemental Figure 1:**
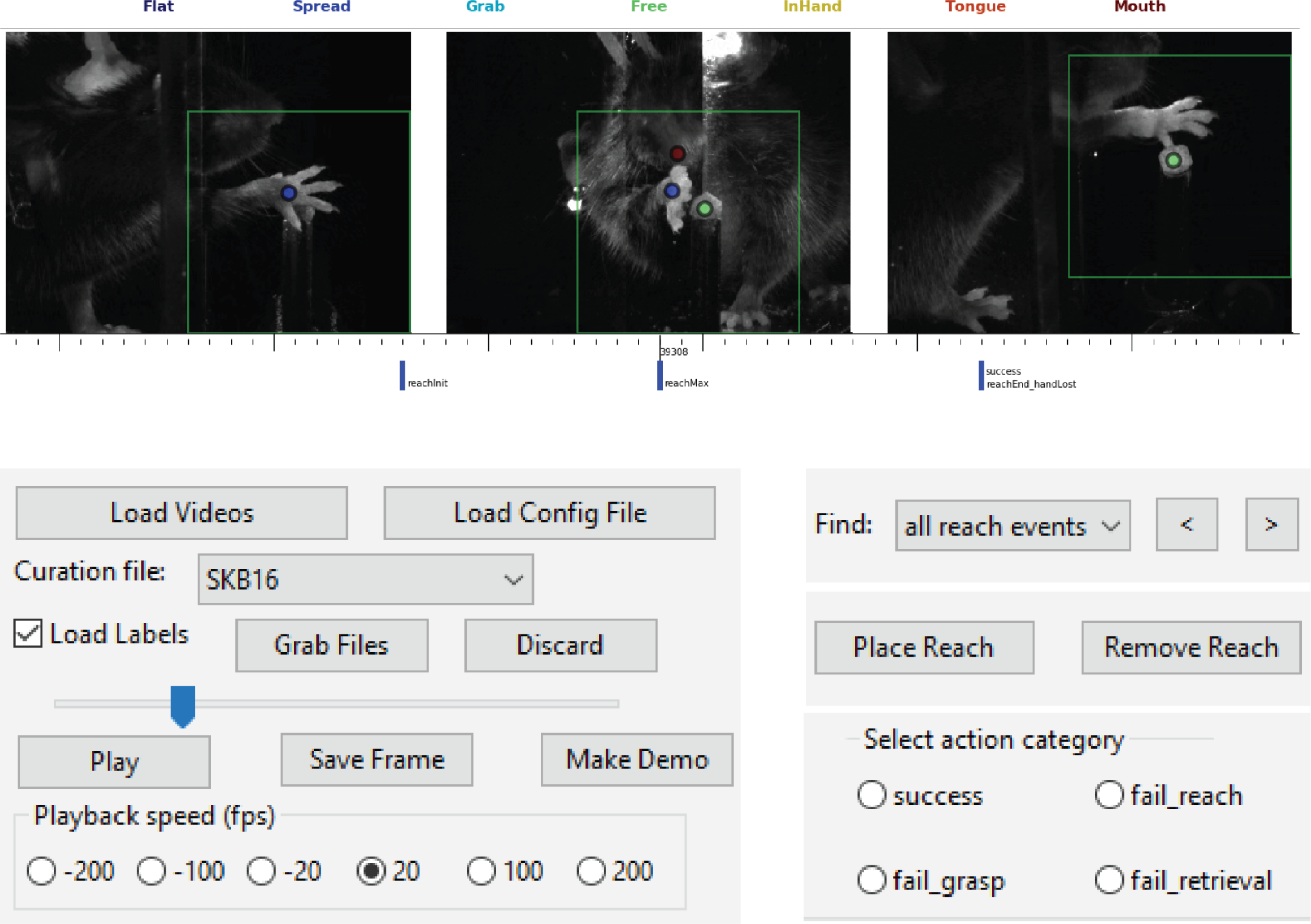
Custom video curation GUI for post hoc analysis. The video curator allows researchers to rapidly navigate video of training sessions using the automatically identified reach features: Reach initiation, Reach max, Reach end and Behavior classification. Researchers can also manually annotate reach outcomes to further subdivide reaches beyond the automatically classified success and failure categories. All timepoints and annotations are used to generate a spreadsheet containing the data from each video session.

**Supplemental figure 2:**
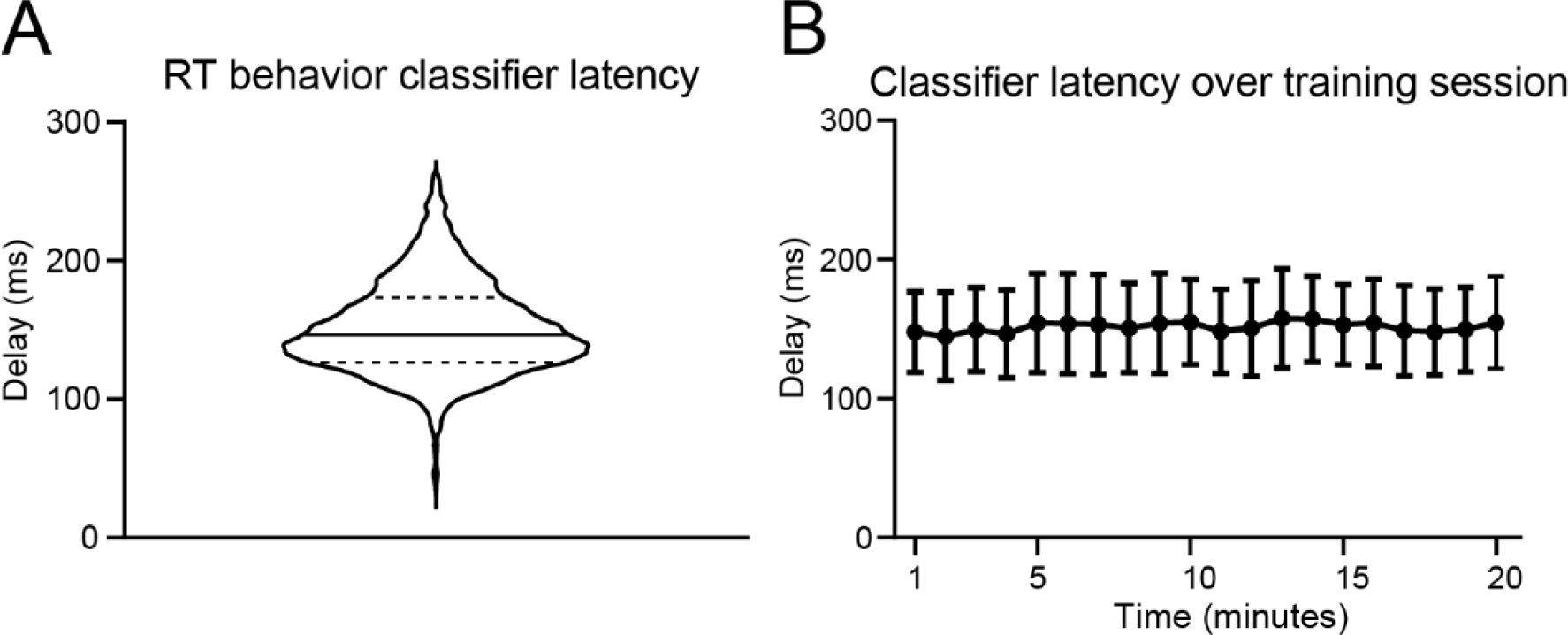
Real-time behavior classifier and hardware latency. A) CLARA classifies reach outcome 151±32ms after the pellet is pulled back into the cage. Dotted lines represent interquartile range. B). The classifier delay remains constant over the course of a training session (20 minutes). Error bars represent SD.

**Supplemental figure 3:**
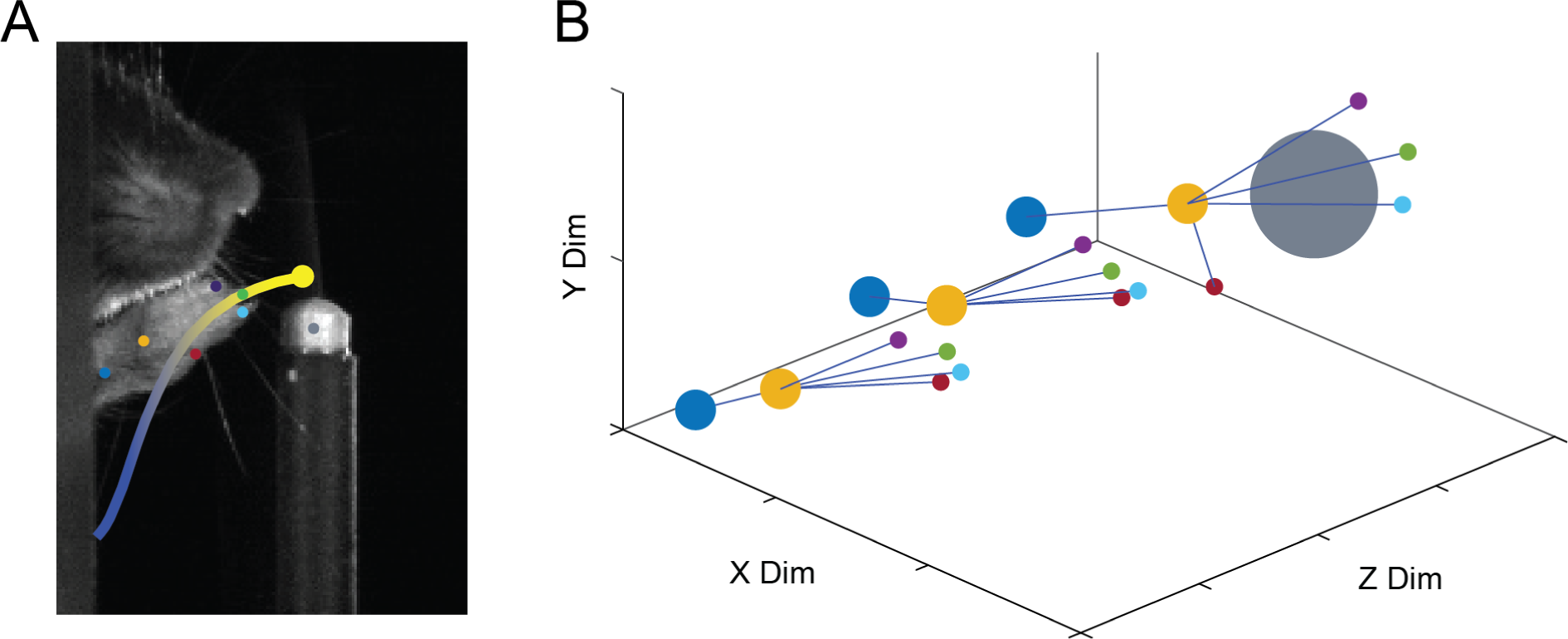
CLARA can track multiple points on the forelimb with single digit resolution. A) Example image of right mouse paw with automatic labels on four fingers, the hand center (orange), wrist (dark blue), and pellet (grey). Line denotes the trajectory of the second finger (green) during the outward phase of the reach. Line color represents normalized time. B) Example of a single reach a three timepoints during the outward phase of the reach plotted in three dimensions.

**Supplemental Figure 4:**
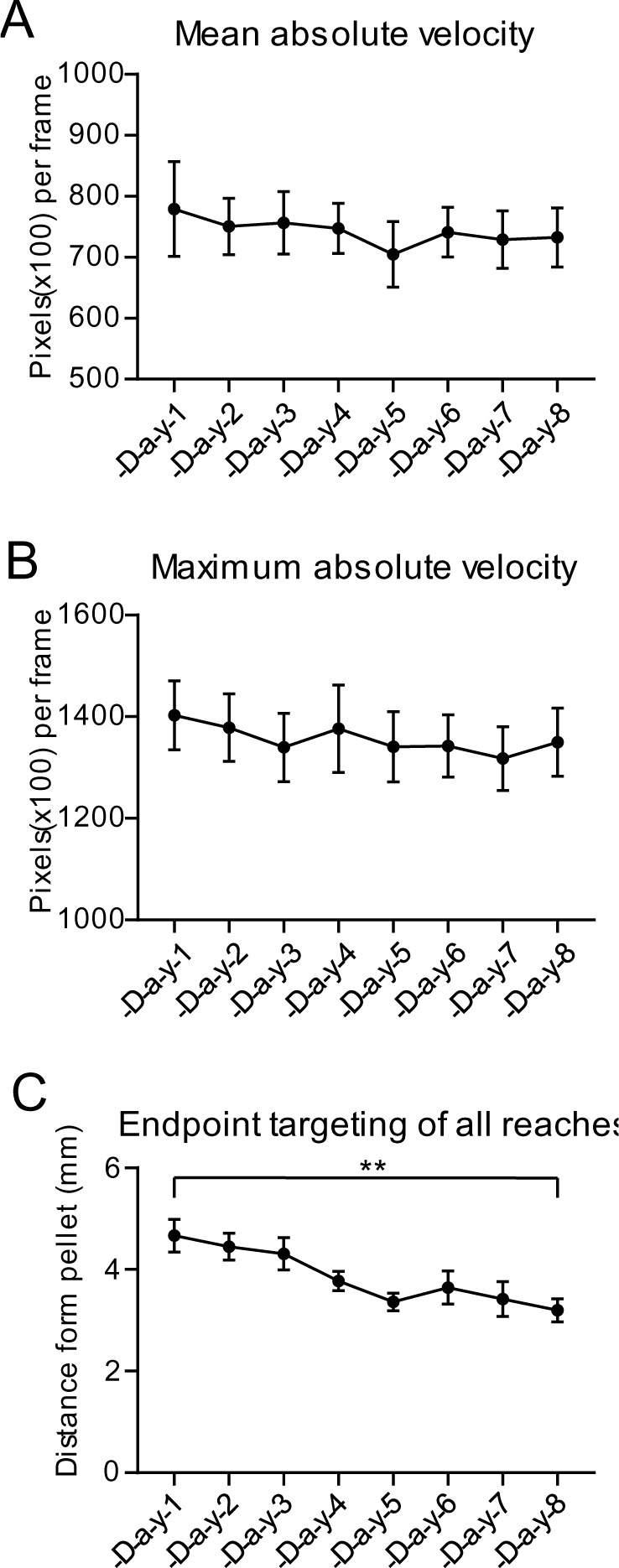
Velocity and targeting data for all reaches over learning. A) Mean absolute velocity during the outward phase of the reach for n=7 across 8 days of learning. C) Maximum absolute velocity during the outward phase of the reach for n=7 across 8 days of learning. Error bars denote standard error. C) Absolute distance of hand center from pellet center at reach max timepoint for n=7 mice. Reaches are targeted closer to the pellet on the day 8 (3.20±0.60mm) of learning compared to day 1 (4.66±0.85mm), p=0.0005 Paired t-test). Error bars denote standard error.

